# Tau in cerebrospinal fluid induces neuronal hyperexcitability and alters hippocampal theta oscillations

**DOI:** 10.1101/2023.01.24.525362

**Authors:** Jessica Brown, Elena Camporesi, Juan Lantero-Rodriguez, Maria Olsson, Alice Wang, Blanca Medem, Henrik Zetterberg, Kaj Blennow, Thomas K. Karikari, Mark Wall, Emily Hill

## Abstract

Alzheimer’s disease (AD) and other tauopathies are characterized by the aggregation of tau into soluble and insoluble forms (including tangles and neuropil threads). In humans, a fraction of both phosphorylated and non-phosphorylated N-terminal to mid-domain tau species, including the aggregated forms, are secreted into cerebrospinal fluid (CSF). Some of these CSF tau species can be measured as diagnostic and prognostic biomarkers, starting from early stages of disease. While in animal models of AD pathology, soluble tau aggregates have been shown to disrupt neuronal function, it is unclear whether the tau species present in CSF will modulate neural activity. Here, we have developed and applied a novel approach to examine the electrophysiological effects of CSF from patients with a tau-positive biomarker profile. The method involves incubation of acutely-isolated wild-type mouse hippocampal brain slices with small volumes of diluted human CSF, followed by a suite of electrophysiological recording methods to evaluate their effects on neuronal function from single cells through to the network level. Comparison of the toxicity profiles of the same CSF samples, with and without immuno-depletion for tau, has enabled a pioneering demonstration that CSF-tau potently modulates neuronal function. We demonstrate that CSF-tau mediates an increase in neuronal excitability in single cells. We then observed, at the network level, increased input-output responses and enhanced paired-pulse facilitation as well as an increase in long-term potentiation. Finally, we show that CSF-tau modifies the generation and maintenance of hippocampal theta oscillations, which have important roles in learning and memory and are known to be altered in AD patients. Together, we describe a novel method for screening human CSF-tau to understand functional effects on neuron and network activity, which could have far-reaching benefits in understanding tau pathology, thus allowing for the development of better targeted treatments for tauopathies in the future.

**Graphic Abstract:** 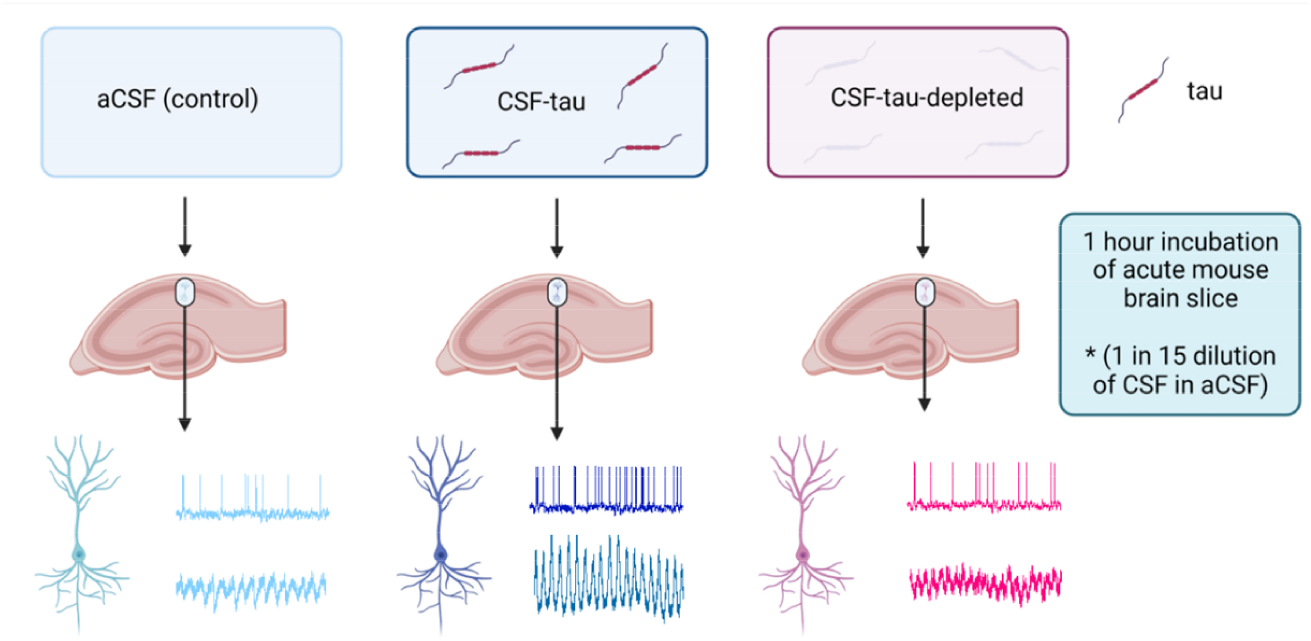

## 1. Introduction

Tau plays a central role in the neuropathology of Alzheimer’s disease (AD) and other tauopathies [67, 110]. Whilst insoluble high-molecular weight tau aggregates, including neurofibrillary tangles are more prominent in late phases of these diseases and can help stage their pathological and clinical severity [20, 29, 92], the soluble forms of tau, including low-molecular weight aggregated and phosphorylated forms, are thought to be the most toxic species that modulate changes in neuronal function in the early stages of disease. In animal models of AD pathology, soluble tau aggregates have been shown to disrupt neuronal function, alter synaptic plasticity and impair cognitive function [19, 43, 79, 102, 105].

In AD,N-terminal to mid-domain tau fragments, both phosphorylated and non-phosphorylated variants, are released into the cerebrospinal fluid (CSF) early in disease progression, which has enabled the development of biomarkers for disease prognosis, diagnosis, and staging [18, 31, 55, 65, 107, 108]. While the presence of different tau forms in the CSF of AD patients is associated with neurofibrillary tangle pathology and cognitive decline [17, 28, 48, 62, 72, 87, 113], it is currently unclear whether the tau present in CSF is functionally active and can therefore modulate neural activity.

We have previously developed a technique using whole-cell patch-clamp recording to introduce structurally-defined soluble full-length recombinant tau aggregates into single neurons [58–60]. Using this approach, we have been able to provide detailed characterisation of the pathological actions of soluble tau aggregates on single neuron electrophysiology, evaluate changes to synaptic transmission and plasticity, and demonstrate that the effects are specific to the aggregates as monomers had little or no effect [58, 59]. However, a limitation of using recombinantly produced tau is that it cannot represent the full range of endogenous tau forms present in the disease and delivery via a patch-pipette will only target a single neuron, thus, it is not possible to determine how these changes will modulate brain network function. To overcome these drawbacks, we have developed a new approach of incubating cortical slices in dilutions of patient CSF, which has allowed us to examine effects of clinically derived tau species on single neurons and on cortical network activity (theta oscillations).

Active cortical networks support several rhythms (such as gamma and theta oscillations) that can be measured clinically including in individuals suspected to be affected by AD. Theta oscillations (4 – 7 Hz) represent the largest extracellular signal recorded in the mammalian brain [124] and are considered integral to hippocampal learning and memory processes [13, 15, 51]. Specifically, activity within the theta band reflects the temporal encoding of information and is thus associated with the consolidation and retrieval of episodic and spatial memory [82, 127]. Work on both rodent models of AD and human AD patients has shown that these rhythms and their coupling are disrupted as the pathology progresses [45, 64, 89, 90, 98, 109, 111]. In humans, changes in theta power have been widely reported during the performance of spatial navigation, working memory and recognition tasks [21, 25, 34, 40, 75, 103].

In this study, we have characterised the functional neurotoxic changes induced by tau-positive CSF samples, on neuronal function. We have used a suite of detailed electrophysiological measures; from single cells to network activity. We demonstrate that CSF-tau mediates an increase in neuronal excitability, alters synaptic transmission and plasticity and modifies the generation and maintenance of hippocampal theta oscillations. However, the same CSF samples, with tau removed by immuno-depletion, failed to produce these effects, indicating that tau is central to these neural changes.

## 2. Materials and methods

### 2.1 Collection, biomarker profiling and immuno-depletion of CSF

All CSF samples were obtained from patients by lumbar puncture from the L3/L4 or the L4/L5 inter-space in the morning, according to standardized procedure. Samples were collected in a propylene tube and centrifuged for 10 min at 1800 x g at + 4 °C. The supernatant was stored at – 80 °C prior to use. The CSF samples used in this study were the remainder following clinical routine analyses, from multiple de-identified patients, pooled together in a procedure approved by the Ethics Committee at the University of Gothenburg (#EPN 140811). The collection and use of CSF samples were in accordance with the Swedish law of biobank in healthcare (2002:297).

The biomarker profile for the pool, including Aβ42, Aβ40, total tau (t-tau) and phosphorylated tau at threonine 181 (p-tau181), was measured with the fully automated Lumipulse (Fujirebio) platform following published protocols [49]. The CSF pool was then divided into two equal aliquots. To remove tau protein, aliquot 1 was immuno-depleted using immunoprecipitation (IP) following a previously established protocol [77] combining three antibodies to thoroughly cover across the full-length tau-441 sequence. Four μg of Tau12 (N-terminal [epitope = amino acids 6-18], BioLegend), HT7 (mid-region [epitope = amino acids 159-163], ThermoFisher), and TauAB (C-terminal [epitope = amino acids 425-441], kindly provided by MedImmune) were independently conjugated with 50 μL of M-280 Dynabeads (sheep anti-mouse IgG, Invitrogen) following manufacturer’s recommendations. Beads were then mixed and added to aliquot 1. Incubation was performed overnight at 4 °C on a rolling station to allow mixing of the beads with the sample and to avoid the beads forming aggregated deposits. The next morning, the beads were removed from the mixture by standing the tube against a magnet. The supernatant was carefully pipetted out to obtain the CSF-tau-depleted fraction. Aliquot 2 underwent the same free-thaw cycle as the CSF-tau-depleted but was not immuno-depleted. Both samples were stored at −80 °C prior to use. To verify the removal of tau after the immuno-depletion, we used different assays that cover various regions of tau. The t-tau assay from Quanterix, p-tau181 [73] and p-tau231 [8] in house assays were used for tau measurements in both the CSF-depleted and the non-depleted aliquots on a Simoa HD-X instrument (Quanterix, Billerica, MA, USA) following methods originally described in the cited publications. To further validate the removal of tau, the CSF-depleted sample was also tested using the mid-region targeting Lumipulse p-tau-181 assay (Fujirebio) [49].

CSF collection and processing were performed at the University of Gothenburg, Sweden, with local ethical approval, and the samples sent to the University of Warwick, UK, where all experiments were done as approved by the local Human Tissue Authority and Biomedical & Scientific Research Ethics Committees.

### 2.4 Preparation of mouse brain slices

All animal care and experimental procedures were reviewed and approved by the institutional animal welfare and ethical review body (AWERB) at the University of Warwick. Animals were kept in standard housing with littermates, provided with food and water ad libitum and maintained on a 12:12 (light-dark) cycle. C57BL/6 mice (3–4-week-old; male and female) were killed by cervical dislocation and decapitated in accordance with the United Kingdom Animals (Scientific Procedures) Act (1986). The brain was rapidly removed, and acute parasagittal or horizontal brain slices (350-400 μm) were cut with a Microm HM 650V microslicer in cold (2–4°C) high Mg^2+^, low Ca^2+^ artificial CSF (aCSF), composed of the following: 127 mM NaCl, 1.9 mM KCl, 8 mM MgCl_2_, 0.5 mM CaCl_2_, 1.2 mM KH_2_PO_4_, 26 mM NaHCO_3_, and 10 mM D-glucose (pH 7.4 when bubbled with 95% O_2_ and 5% CO_2_, 300 mOsm). Slices were stored at 34°C in standard aCSF (1 mM Mg^2+^ and 2 mM Ca^2+^) for between 1 and 8 hours.

### 2.5 Incubation of acute brain slices with CSF samples

After at least 1 hour of recovery, slices were either incubated in aCSF (control) or in varying dilutions (1:15, 1:30, 1:100 in aCSF) of CSF-tau or CSF-tau-depleted (immunodepleted for tau) for 1 hour, in bespoke incubation chambers at room temperature. The incubation chambers consisted of small, raised grids (to allow perfusion of slices from above and below) placed in the wells of a twenty-four well plate (Falcon) which were bubbled with 95 % O_2_, 5 % CO_2_ using microloaders (Eppendorf). Slices were placed into the incubation chambers one at a time (minimum volume 1.5 ml to cover raised grid). Individual slices were then placed on the recording rig and perfused with regular aCSF throughout the recording period, so the CSF-tau was only present for the 1-hour incubation.

### 2.6 Whole cell patch clamp recording from single hippocampal CA1 pyramidal neurons

A slice was transferred to the recording chamber, submerged, and perfused (2–3 ml/min) with aCSF at 30°C. Slices were visualised using IR-DIC optics with an Olympus BX151W microscope (Scientifica) and a CCD camera (Hitachi). Whole-cell current-clamp recordings were made from pyramidal cells in area CA1 of the hippocampus using patch pipettes (5–10 MΩ) manufactured from thick-walled glass (Harvard Apparatus). Pyramidal cells were identified by their position in the slice, morphology (from fluorescence imaging) and characteristics of the standard step current-voltage relationship. Voltage recordings were made using an Axon Multiclamp 700B amplifier (Molecular Devices) and digitised at 20 kHz. Data acquisition and analysis were performed using pClamp 10 (Molecular Devices). Recordings from neurons that had a resting membrane potential of between −55 and −75 mV at whole-cell breakthrough were accepted for analysis. The bridge balance was monitored throughout the experiments and any recordings where it changed by >20% were discarded.

#### Stimulation protocols

To extract the electrophysiological properties of recorded neurons, both step, ramp and more naturalistic, fluctuating currents were injected.

#### Standard IV protocol

The standard current-voltage relationship was constructed by injecting standard (step) currents from – 200 pA incrementing by either 50 or 100 pA (1 s duration) until a regular firing pattern was induced. A plot of step current against voltage response around the resting potential was used to measure the input resistance (from the gradient of the fitted line).

#### Rheobase ramp protocol

To evaluate the rheobase (the minimum current required to elicit an action potential (AP)) a current ramp was injected into neurons. From the baseline, a 100 ms duration ramp to −100 pA was followed by a 900 ms duration ramp at 0.33 pA/ms up to 200 pA, then a step back down to baseline (zero current).

#### Dynamic IV protocol

The dynamic-I-V curve, defined by the average transmembrane current as a function of voltage during naturalistic activity, can be used to efficiently parameterize neurons and generate reduced neural models that accurately mimic the cellular response. The method has been previously described [10, 11, 56], for the dynamic-IV computational code, see [56]. Briefly, a current wave form, designed to provoke naturalistic fluctuating voltages, was constructed using the summed numerical output of two Ornstein–Uhlenbeck processes [121] with time constants τfast = 3 ms and τslow = 10 ms. This current waveform, which mimics the stochastic actions of AMPA and GABA-receptor channel activation, is injected into cells and the resulting voltage recorded (a fluctuating, naturalistic trace). The voltage trace was used to measure the frequency of action potential firing and to construct a dynamic-I-V curve. The firing rate was measured from voltage traces evoked by injecting a current wave form of the same gain for all recordings (to give a firing rate of ~2–3 Hz). Action potentials were detected by a manually set threshold and the interval between action potentials measured. Dynamic I-V curves were constructed and used to extract a number of parameters including the capacitance, time constant, input resistance, resting membrane potential, spike threshold and spike onset (Fig. 2; [10, 11]). Using these parameters in a refractory exponential integrate-and-fire (rEIF) model reliably mimics the experimental data, with a spike prediction of ~70–80% as shown previously [10, 11]. All analyses of the dynamic-I-V traces were completed using either MATLAB or Julia software platforms [16].

**Figure 1.**
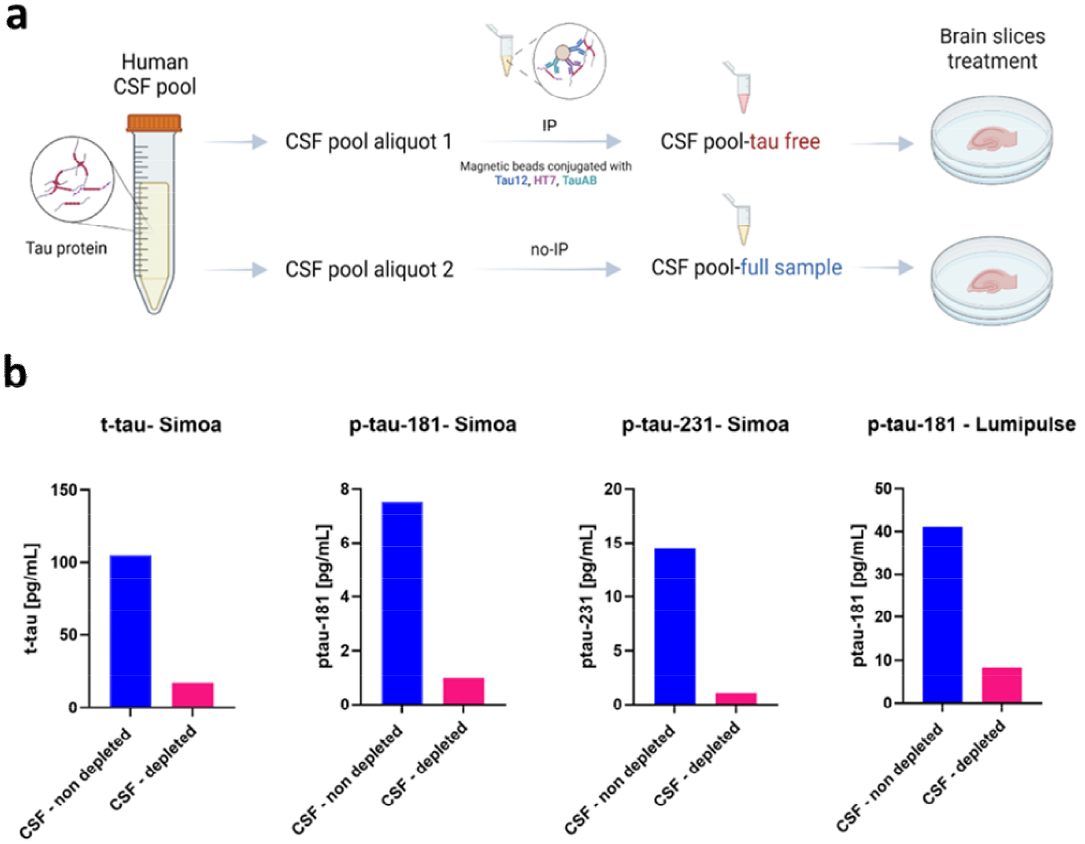
Immunodepletion of pooled CSF to remove tau using a combination of antibodies targeting the N terminus, mid region, and C terminus of tau. **a.** An aliquot of the pooled CSF-tau sample was immuno-depleted for tau by the simultaneous application of the monoclonal antibodies Tau12, HT7 and TauAB (epitopes: amino acids 6-18, 159-163 and 425-441, respectively) that together cover nearly the entire tau-441 protein sequence using published protocols [77]. **b**, To verify the removal of tau after the immuno-depletion, t-tau assay from Quanterix, p-tau181 [73] and p-tau23l [8] in house assays were used for tau measurements in both the CSF-depleted and non-depleted aliquots on a Simoa HD-X instrument (Quanterix, Billerica, MA, USA) following methods originally described in the cited publications and validated using fully automated Lumipulse (Fujirebio) platform following published protocols [49].

**Figure 2.**
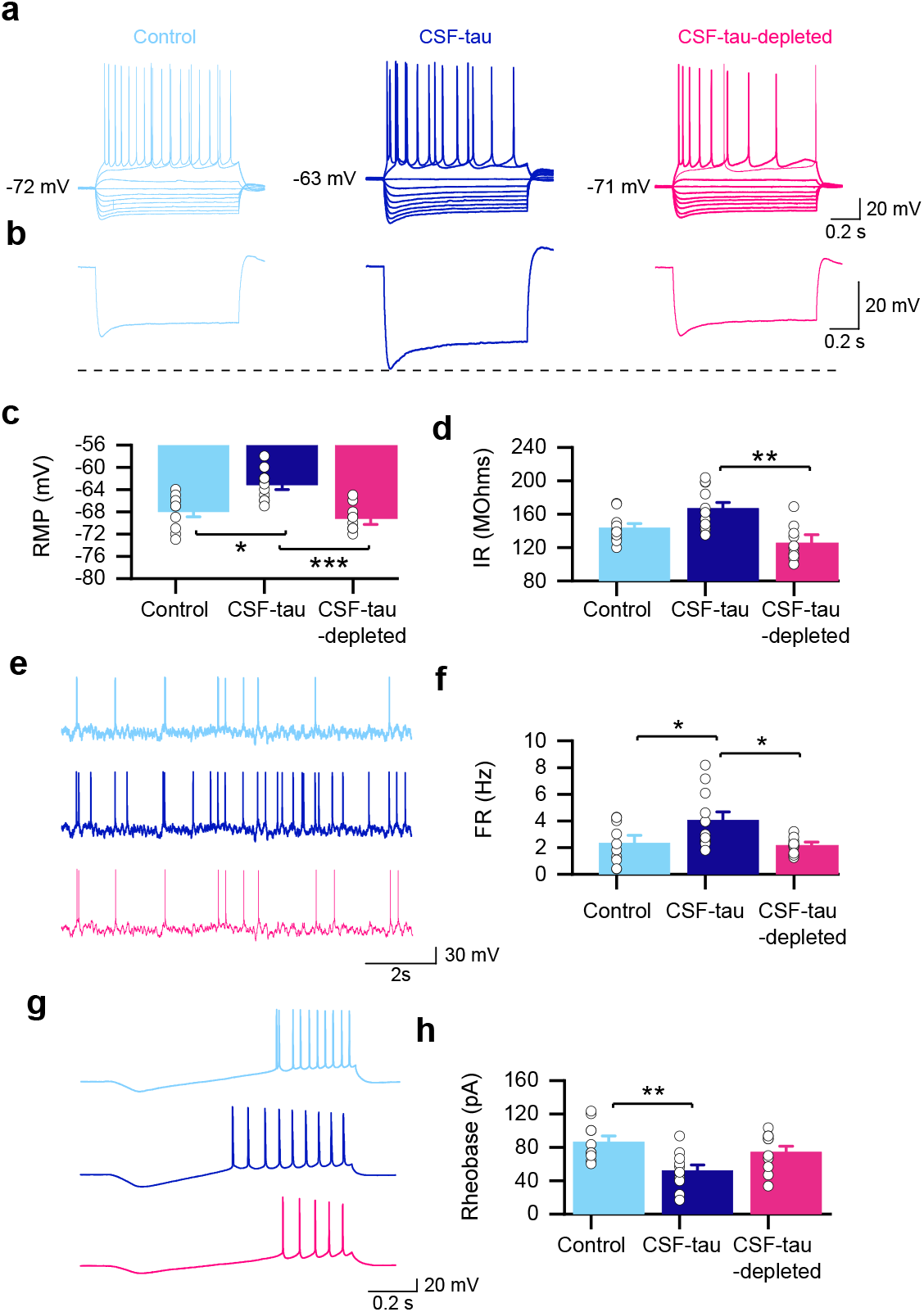
CSF-tau enhances hippocampal pyramidal neuronal excitability. **a.** Representative examples of standard current-voltage responses for slices that have been incubated in control aCSF (light blue; n=10), CSF-tau (CSF; dark blue; n=11) or CSF-tau immunodepleted for tau (CSF-depleted; Pink; n=10). See Materials and Methods for incubation protocols. CSF-tau depolarised the resting membrane potential of the recorded neurons. **b**, the most negative step from traces in (**a**), clearly highlighting that CSF-tau also mediates an increase in input resistance. **c**, CSF-tau incubation resulted in a significant depolarisation of the resting membrane potential (p < 0.0006), which was not observed in the CSF sample immuno-depleted for tau. **d.** CSF-tau incubation also significantly increased input resistance (p < 0.0020), an effect which was also not observed in the CSF sample that was immunodepleted for tau. **e.** Representative example of membrane-potential responses to naturalistic current injection for each of the three conditions. **f.** CSF-tau significantly increased the firing rate (a correlate of neuronal excitability; p < 0.0055). No change was observed with the CSF-tau sample immunodepleted for tau compared to control. **g.** The rheobase current (minimal current to evoke an AP) was determined by injecting a current ramp (−50 to 200 pA) and measuring the minimum current required to fire an action potential. **h.** CSF-tau incubation significantly decreased the rheobase (p = 0.0093). *Panels a, d and f show representative example traces and b, c, e, and g show the mean data and SEM, with individual datapoints overlaid*.

### 2.7 Recording and analysing miniature excitatory postsynaptic currents (mEPSCs)

AMPA receptor-mediated miniature excitatory postsynaptic currents (mEPSCs) were recorded as previously described [37]. To isolate AMPA-mediated mEPSCs, aCSF contained 1 μM TTX, 50 μM picrotoxin to block GABA_A_ receptors and 5 μM L689,560 to block NMDA receptors. Recordings of mEPSCs were obtained at a holding potential of −60 mV using an Axon Multiclamp 700B amplifier (Molecular Devices), filtered at 3 kHz and digitised at 20 kHz (Digidata 1440A, Molecular Devices). Data acquisition was performed using pClamp 10 (Molecular Devices). Analysis of mEPSCs was performed using MiniAnalysis software (SynaptoSoft). Events were manually analysed and were accepted if they had amplitude >6 pA (events below this amplitude were difficult to distinguish from baseline noise) and had a faster rise than decay. Statistical significance was measured using a one-way ANOVA.

### 2.8 Extracellular recording of synaptic transmission and plasticity

A 400 μM parasagittal slice was transferred to the submerged recording chamber and perfused with aCSF at 4-6 ml/min (32°C). The slice was placed on a grid allowing perfusion above and below the tissue and all tubing (Tygon) was gas tight (to prevent loss of oxygen). To record field excitatory postsynaptic potentials (fEPSPs), an aCSF-filled microelectrode was placed on the surface of stratum radiatum in CA1. A bipolar concentric stimulating electrode (FHC) controlled by an isolated pulse stimulator model 2100 (AM Systems, WA) was used to evoke fEPSPs at the Schaffer collateral–commissural pathway. Field EPSPs were evoked every 30 s (0.03 Hz). Stimulus input/output curves for fEPSPs were generated using stimulus strength of 1-5 V for all slices (stimulus duration 200 μs). For the synaptic plasticity experiments, the stimulus strength was set to produce a fEPSP slope ~ 40 % of the maximum response and a 20-minute baseline was recorded before plasticity induction. Paired-pulse facilitation was measured over an interval range of 20 to 500 ms. Long-term potentiation (LTP) was induced by high frequency stimulation (HFS, 100 stimuli in 1s, 100 Hz) and then fEPSPs were recorded for 60 minutes following LTP induction. Signals were filtered at 3 kHz and digitised on-line (10 kHz) with a Micro CED (Mark 2) interface controlled by Spike software (Vs 6.1, Cambridge Electronic Design, Cambridge UK). The fEPSP slope was measured from a 1 ms linear region following the fibre volley.

### 2.9 Theta Oscillations

#### 2.9.1 Induction of theta oscillations in CA3

A 400 μM horizontal slice was transferred to an interface recording chamber (Digitimer BSC3) and placed on a mesh support at the interface of an oxygen-rich atmosphere and underlying aCSF. The temperature of the aCSF was set at 31°C and the flow rate was 1.5 ml/min. An aCSF-filled glass microelectrode was placed in CA3 to record activity. Baseline activity was recorded for 20 minutes before the bath application of the acetylcholine receptor agonist carbachol, selected to induce oscillations given the evidenced role of acetylcholine receptors in regulating hippocampal-dependent oscillations [7, 119]. Carbachol was applied at 50 μM, a concentration previously demonstrated to induce CA3 theta activity in rodent slices [3, 44, 70, 126]. Theta oscillations were recorded for 30 minutes with a differential amplifier (Warner instrument DP-301) with filter settings: low pass 3KHz, high pass 1Hz, amplification x1000). Data was acquired at a sampling rate of 10 KHz with a Micro1401 (Cambridge Electronic Design) using Spike 2 software.

#### 2.9.2 Analysis of theta oscillations

Carbachol-elicited oscillations were characterised using power spectral density (PSD) analysis in Spike 2. PSD profiles of the field potential recordings filtered for the theta band (4 – 7 Hz) were generated by Fourier transform analysis (Hanning window, FFT size 2048, resolution 4.883 Hz). The profiles were calculated from a 100 – 300 second section of the field potential trace displaying peak oscillatory activity, with the baseline power subtracted offline (supplementary figure 2). Recordings from slices from each treatment condition (control, CSF-tau and CSF-tau-depleted) were conducted in a random order each day to prevent any time effects, and a slice was only included in the analysis if robust oscillations were evoked. Differences between the three treatment groups (control, n = 8; CSF-tau, n = 8; CSF-tau-depleted, n = 8) in mean oscillatory power (mV^2^) were compared using a Kruskal-Wallis test followed by a Dunn’s multiple comparisons test. The peak oscillatory power of each trace (n=8 per group) was also compared with the corresponding baseline (n = 8 per group) within each treatment group and mean differences assessed using Wilcoxon tests. Finally, the mean time period (latency, seconds) from application of carbachol to the onset of oscillatory activity was compared between treatment groups using a Kruskal-Wallis test followed by a Dunn’s multiple comparisons test.

### 2.10 Drugs

Picrotoxin (PTX; Sigma), carbachol (carbamoylcholine chloride; Sigma), Tetrodotoxin (TTX; Sigma) and trans-2-carboxy-5,7-dichloro-4-phenylaminocarbonylamino-1,2,3,4-tetrahydroquinoline (L689,560; Hello-Bio) were made up as stock solutions (1-50 mM) in distilled water or dimethyl sulfoxide (DMSO) and then diluted in aCSF on the day of use. The concentration of DMSO did not exceed 0.1 % in the final solutions.

### 2.11 Statistical Analysis

Statistical analysis was performed using GraphPad Prism. Due to small sample sizes (n < 15), statistical analysis was performed using non-parametric methods; Kruskal–Wallis analysis of variance (ANOVA), Mann–Whitney and Wilcoxon signed-rank tests as required. All data are represented as mean and standard error of the mean with individual experiments represented by single data points. For all experiments, significance was set at p ≤ 0.05. Data points for each experimental condition were derived from a minimum of four individual animals.

## 3. Result

### 3.1 Characterisation of CSF-tau before and after the immuno-depletion for tau

To investigate the effects of CSF-tau on neuronal properties, in this proof-of-principle study, we used pooled CSF from multiple de-identified patients. The biomarker profile for the pool was measured with the Lumipulse platform. For the pooled samples used in this study, CSF Aβ1-42 level (469 pg/ml) was abnormal as it was below the clinically validated abnormality threshold of 526 pg/ml [48]. Aβ1-42/Aβ1-40 ratio was also in the positivity range (<0.072 pg/ml). T-tau (measured = 542 pg/ml) was above the abnormality threshold of 409 pg/ml [80] but p-tau181 remained negative [48]. These profiles suggest that pooled CSF sample was amyloid-positive, phosphorylated-tau-negative but neurodegeneration-positive (A+T-N+) [67].

To allow determination of the specific effects of tau on electrophysiological properties, an aliquot of the pooled CSF-tau sample was immuno-depleted for tau by the simultaneous application of the monoclonal antibodies Tau12, HT7 and TauAB (epitopes: amino acids 6-18, 159-163 and 425-441 respectively) that together cover a wide span of the tau-441 protein sequence. To confirm that tau removal protocols were effective, the immuno-depleted samples were reanalysed for t-tau, p-tau181 and ptau-231 using Simoa assays. P-tau181, p-tau231 and total-tau signals decreased by 86.9% and 92.4% and 83.8%, respectively. To further validate the depletion, p-tau181 was measured using a fully automated Lumipulse platform and the signal decreased by 80.5% (figure 1).

### 3.2 CSF-tau enhances hippocampal pyramidal neuronal excitability

Tau aggregates have previously been reported both *in vitro* and *in vivo* to affect neuronal function, excitability, and synaptic plasticity [19, 43, 59, 79, 102, 105]. These tau-mediated deficits are largely mediated by soluble forms of the protein [59, 93, 112, 125], including those isolated from AD brains [78, 79]. However, studies using patient-derived CSF samples, which are estimated to contain a fraction of the soluble tau aggregates from AD brains [71, 87, 107], are lacking. Here, we investigated if the reported pathological effects of soluble tau aggregates can be reproduced with CSF-tau.

In this study, we used pooled CSF samples from multiple patients. The relatively large volume of pooled sample (5 ml) allowed us to perform a wide range of experiments using the same sample to fully characterise its neurotoxic effects. We diluted the CSF-tau sample with aCSF to reduce the volume of sample that was used for each experiment. As our eventual end goal will be to use individual patient samples (much smaller volumes ~300 μl) to enable correlation with clinical data, we first sought to find the minimum volume (maximum dilution in aCSF) of CSF-tau, for which we could detect a change in neuronal function. We tested 1:100, 1:30 and 1:15 dilutions of the CSF-tau in aCSF. We used neuronal excitability, measured using whole-cell patch clamp recording from single CA1 pyramidal cells in the hippocampus, as the functional readout. We have previously found this to be a reliable and robust change induced by recombinant tau [58, 59]. We found no significant changes to neuronal excitability with either 1:100 dilution or 1:30 dilution of CSF-tau, but a significant depolarisation and increase in firing rate was observed with a 1:15 dilution (100 μl CSF in 1.5 ml aCSF; supplementary Figure 1). We therefore used a 1:15 dilution of CSF-tau and CSF-tau-depleted for all subsequent experiments. To fully evaluate the effects of CSF-tau at a single neuron level, slices were either incubated with CSF-tau (n = 11), CSF-tau-depleted (n = 10) or control aCSF (n = 10; no human sample) for 1 hour before recordings were made. Initially, standard current step protocols were used to measure passive neuronal parameters (see materials and methods for details).

Incubation with CSF-tau led to significant depolarisation of the resting membrane potential (RMP; mean RMP in control slices of −68 ± 0.92 mV compared to the mean RMP in CSF-tau of −63 ± 0.79 mV; *Figure 2a, b*), an effect that was not observed in slices incubated in CSF-tau-depleted (mean RMP of −69 ± 1.09 mV; *Figure 2a, b*). A Kruskal-Wallis test showed that the difference in RMP between the three treatment groups was significant *(Figure 2b;* H (2) = 14.98, p < 0.0006), with Dunn’s tests revealing a significant depolarisation of the RMP following CSF-tau treatment relative to aCSF control slices (p < 0.0121) and treatment with CSF-tau-depleted (p < 0.0008), whilst there was no significant difference between control and CSF-tau-depleted groups (p >0.9999).

Incubation with CSF-tau also significantly increased the input resistance (IR, reflecting a decrease in whole cell conductance). The mean IR in control slices was 143.6 ± 5.2 MΩ, compared to the mean IR in slices incubated in CSF-tau of 167 ± 7.2 MΩ; *Figure 2a, c*), an effect which again was not observed in slices incubated in CSF-tau-depleted (mean IR of 126 ± 9.9 mV; *Figure 2a, c*). The difference in IR between the three treatment groups was also significant (Kruskal Wallis test; *Figure 2c;* H (2) = 12.39, p < 0.0020), with Dunn’s tests revealing a significant increase in IR following CSF-tau treatment relative treatment with CSF-tau-depleted (p < 0.0013), whilst there was no significant difference between either control (aCSF) and CSF-tau-depleted groups (p < 0.2831).

Given that CSF-tau depolarises the resting membrane potential and increases the IR, we predicted that it should also increase the action potential firing rate. To test this, we used a 40 s fluctuating naturalistic current injection (to mimic synaptic activation) and measured the firing rate. Incubation with CSF-tau significantly increased firing rate (FR). The mean FR in control was 2.3 ± 0.5 Hz compared to the mean FR in neurons exposed to CSF-tau of 4.2 ± 0.6 Hz; *Figure 2d, e*), This increase in FR was not observed in slices incubated in CSF-tau-depleted (mean FR of 2.1 ± 0.3 Hz; *Figure 2d, e*). A Kruskal-Wallis test showed that the difference in FR between the three treatment groups was also significant *(Figure 2e;* H (2) = 10.4, p<0.0055), with Dunn’s tests revealing a significant increase in FR following CSF-tau treatment relative to aCSF control slices (p < 0.0129) and treatment with CSF-tau-depleted (p < 0.0222), whilst there was no significant difference between control and CSF-tau-depleted groups (p > 0.9999).

We have previously shown that tau aggregates decrease the rheobase (minimum current needed to fire an action potential; [58]), therefore we next checked whether the increase in excitability was accompanied by a change in rheobase (Figure 2f). The mean rheobase in neurons exposed to CSF-tau was significantly decreased (52 ± 6.6 pA) compared to aCSF control slices (87 ± 6.8 pA) or slices incubated in CSF-tau-depleted (74 ± 6.9 pA). A Kruskal-Wallis test showed that the difference in FR between the three treatment groups was also significant *(Figure 2g; H (2)* = 9.3, p < 0.0093), with Dunn’s tests revealing a significant decrease in rheobase following CSF-tau treatment relative to aCSF control slices (p = 0.0080), whilst there was no significant difference between aCSF control and CSF-tau-depleted groups (p = 0.8856).

To determine if other neuronal parameters were significantly altered by CSF-tau, we used the dynamic IV method [10, 11] to measure capacitance, the time constant, the spike threshold and spike onset. Unlike our previous studies, using recombinant tau oligomers and associated truncations [58, 59], there were no significant differences in any of these parameters. We also evaluated whether there were changes to action potential amplitude or half-width, which we have previously observed with recombinant soluble tau aggregates [58, 59]. Again, we observed no significant changes to either of these parameters between the three conditions.

Thus, CSF-tau replicates a number of the pathological effects which have been reported previously in models of tauopathy. These effects were absent when tau was removed by immunodepletion, indicating that tau is critical to these disease-associated changes.

### 3.3 CSF-tau enhanced basal EPSP amplitude and paired pulse facilitation and long-term potentiation

Next, we investigated whether the changes to neuronal excitability in single pyramidal cells were accompanied by changes in synaptic transmission and plasticity in the CA1 region of the hippocampus.

Incubation in CSF-tau led to increased basal synaptic transmission, as measured by stimulus input/output curves (*Figure 3a*). For example, the mean fEPSP slope at 3 V in CSF-tau treated slices was 0.46 ± 0.06 mV/ms (n=8) compared to control slices which was 0.25 ± 0.04 mV/ms (n=10) and 0.22 ± 0.03 mV/ms in CSF-tau-depleted (n=12). A two-way ANOVA evaluating the fEPSP response to increasing stimulus across the three conditions showed a significant difference between the groups (F (25, 225) = 15.05, p < 0.0001) with significantly greater fEPSP slope for CSF-tau compared to CSF-tau-depleted at 2.5 mV (p = 0.0486), 3 mV (p = 0.0241) and 3.5 mV (p = 0.0494) stimulus strengths. We also measured the fibre volley for a given stimulus (4 mV) it found that it was significantly larger in CSF-tau (0.29 ± 0.03 mV) compared to control (0.15 ± 0.02 mV, p=0.0278)

**Figure 3.**
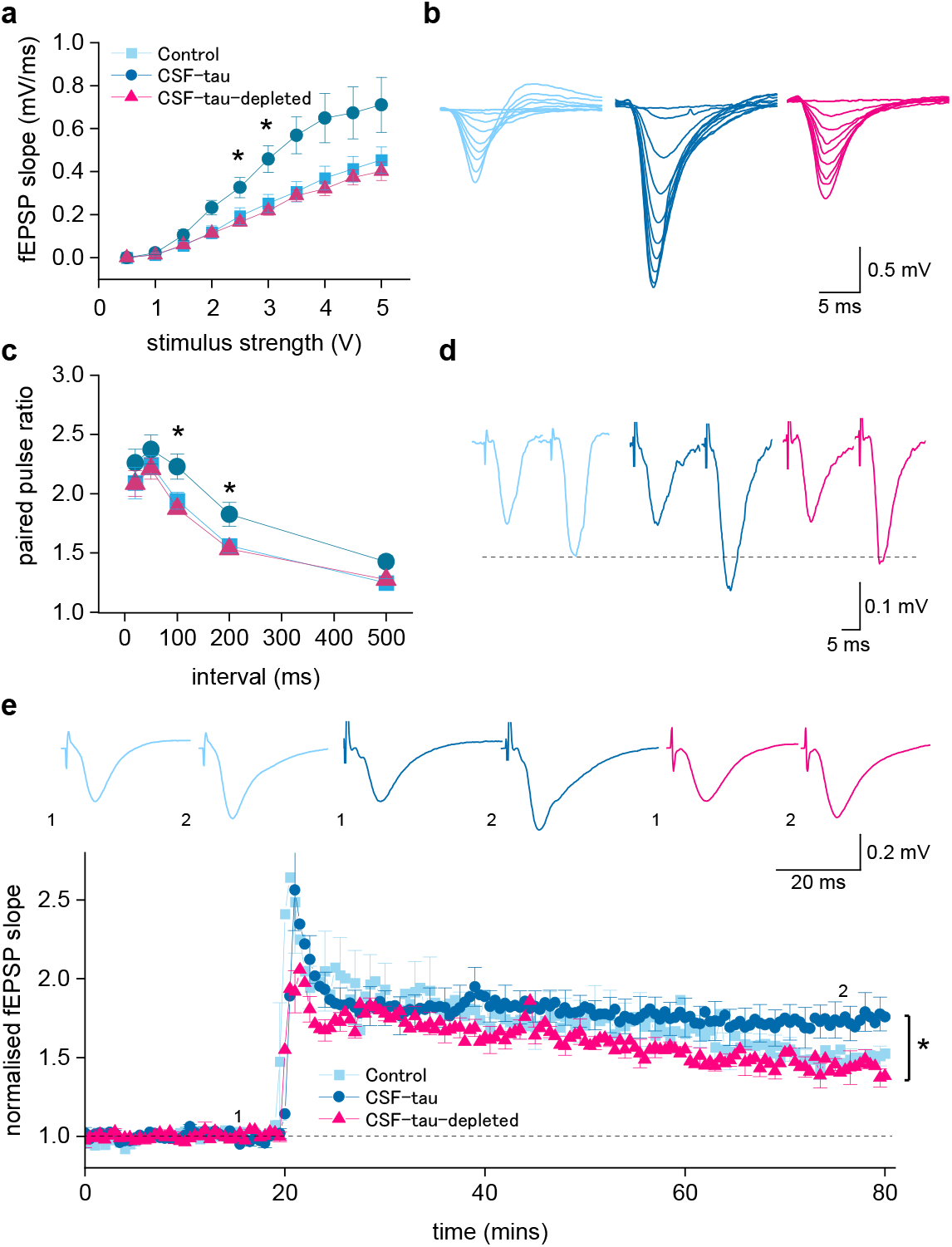
CSF-tau enhances basal synaptic transmission and synaptic plasticity. **a.** Graph plotting mean fEPSP slope against stimulus strength for control (n=10 slices), CSF-tau (n=11 slices) and CSF-tau-depleted (n=10 slices). **b**, Superimposed fEPSP waveforms at increasing stimulus strengths (0.5 to 5 V) for each of the three conditions. CSF-tau significantly enhanced fEPSP slope. **c,** Graph plotting mean paired pulse ratio against interval for all three conditions. CSF-tau significantly enhanced paired pulse facilitation. **d**, Representative example traces of fEPSP waveforms for the 200 ms interval for all three conditions. The first fEPSPs have been normalised so that facilitation can be compared across conditions **e,** Graph plotting mean normalised (to the baseline) fEPSP slope against time for all three conditions. After a 20-minute baseline, LTP was induced by HFS. Inset, example fEPSP waveforms before and after LTP induction (average of waveforms at 75-80 minutes). The mean potentiation was enhanced in CSF-tau. All Data is represented as Mean ± SEM.

We then examined whether there were changes to paired pulse facilitation (PPF). If the increase in fEPSP amplitude is a consequence of an increase in release probability, then it would be predicted that the degree of PPF would be reduced. However, the opposite was observed, and the degree of facilitation was significantly increased at 100 ms and 200 ms intervals (*Figure 3b*). For example, the mean paired-pulse ratio at a 100 ms interval in CSF-tau was 2.23 ± 0.1 compared to control aCSF which was 1.94 ± 0.07 and 1.87 ± 0.04 in CSF-tau-depleted. A two-way ANOVA evaluating the paired-pulse ratio at different intervals across the three conditions showed a significant difference between the groups (F (2, 135) = 10.62, p< 0.0001) with significantly greater facilitation for CSF-tau compared to CSF-tau-depleted at 100 (p = 0.0081) and 200 ms time intervals (p = 0.0368).

In the same recordings, we then examined whether CSF-tau altered long-term synaptic plasticity by recording long term potentiation (LTP) at the CA1 Schaffer collateral synapses. We measured the potentiation 60 minutes after HFS [61, 133] and found LTP to be significantly enhanced by CSF-tau. The mean potentiation (displayed as normalised fEPSP slope, *Figure 3c*) at 55-60 minutes after induction was 1.76 ± 0.09 in CSF-tau, in control aCSF it was 1.51 ± 0.06 and in CSF-tau-depleted it was 1.46 ± 0.07. A Kruskal Wallis test showed that the difference in potentiation between the three treatment groups was significant (H (2) = 6.683, p = 0.0354) with Dunn’s tests revealing a significant increase in potentiation following CSF-tau treatment relative to treatment with CSF-tau-depleted (p = 0.0415), whilst there was no significant difference between control and CSF-tau-depleted groups (p > 0.9999).

### 3.4 Incubation with CSF-tau leads to an increase in the amplitude and frequency of mEPSCs

Since the increase in fEPSP amplitude did not appear to result from an increase in release probability, we hypothesised that it could result from an increase in synaptic AMPA receptor density. To evaluate whether there are changes in postsynaptic AMPA receptor currents, we used whole-cell voltage clamp recordings from CA1 pyramidal neurons to record AMPA receptor-mediated miniature EPSCs (mEPSCs) in the presence of PTX (to block GABA_A_ receptors), L689,560 (to block NMDA receptors) and TTX (to block voltage gated Na channels).

We observed a significant increase in the frequency of mEPSCs (shorter interval between events) in the slices incubated with CSF-tau compared to control slices *(Figure 4a*). The distribution of intervals between events in CSF-tau was significantly different to that of control slices (Kolmogorov-Smirnov test; D = 0.5494, p < 0.0001, *Figure 4b)* with mEPSCs occurring at a much higher frequency in CSF-tau treated slices. On average the frequency was increased 7.5-fold compared to control recordings. Then, by comparing the mean mEPSC intervals between events for each slice across the two conditions, we further validated that slices incubated in CSF-tau showed significantly decreased intervals between events (increased frequency) compared to control slices (mean interval in CSF-tau was 1.9 ± 0.09 s vs mean interval in control was 17.6 ± 1.04 s; Mann-Whitney test p=0.0004; *Figure 4c).*

**Figure 4:**
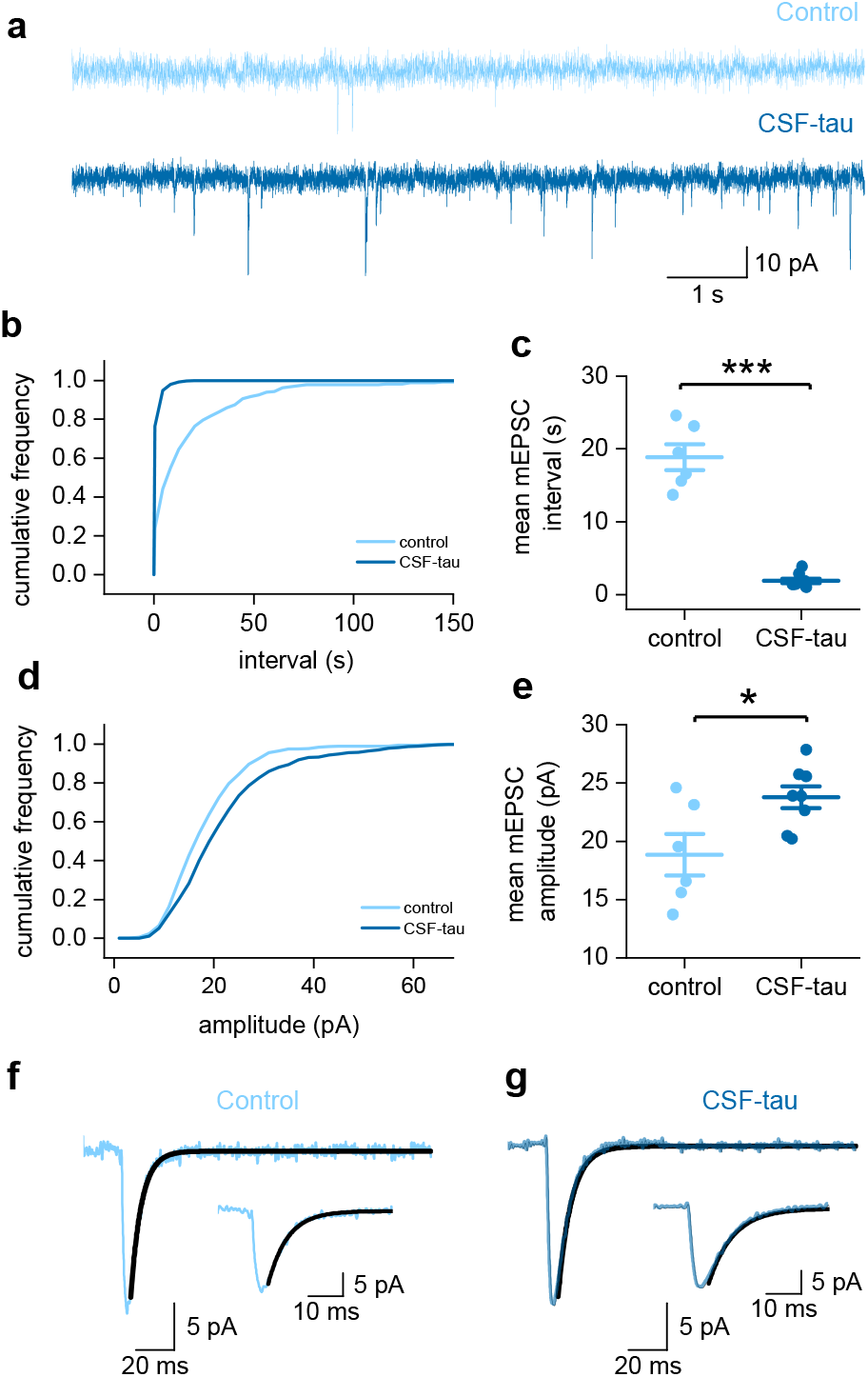
CSF-tau increases the frequency and amplitude of AMPA receptor-mediated mEPSCs. **a,** Representative traces (10 s duration) from control and CSF-tau incubated slices, demonstrating the significant difference in mEPSC frequency. **b**, Cumulative frequency plot of mEPSC interval (between events). **c,** Mean mEPSC interval for each slice is plotted. CSF-tau decreases the interval between mEPSCs, representing an increase in frequency, compared to control (P=0.0004). **d**, Cumulative frequency plot of mEPSC amplitude. **e**, Mean mEPSC amplitude for each slice is plotted. CSF-tau significantly enhances the amplitude of mEPSCs compared to control (p=0.0426). **f, g**, The average mEPSC waveforms for each recording were analysed for kinetics (10-90% for the rise time and exponential fit for the decay time). An example is shown for control (f) and CSF-tau (g), and the exponential fit is overlaid in black. Inset, higher magnification to show the decay fit. Data is represented as Mean ± SEM.

**Figure 5.**
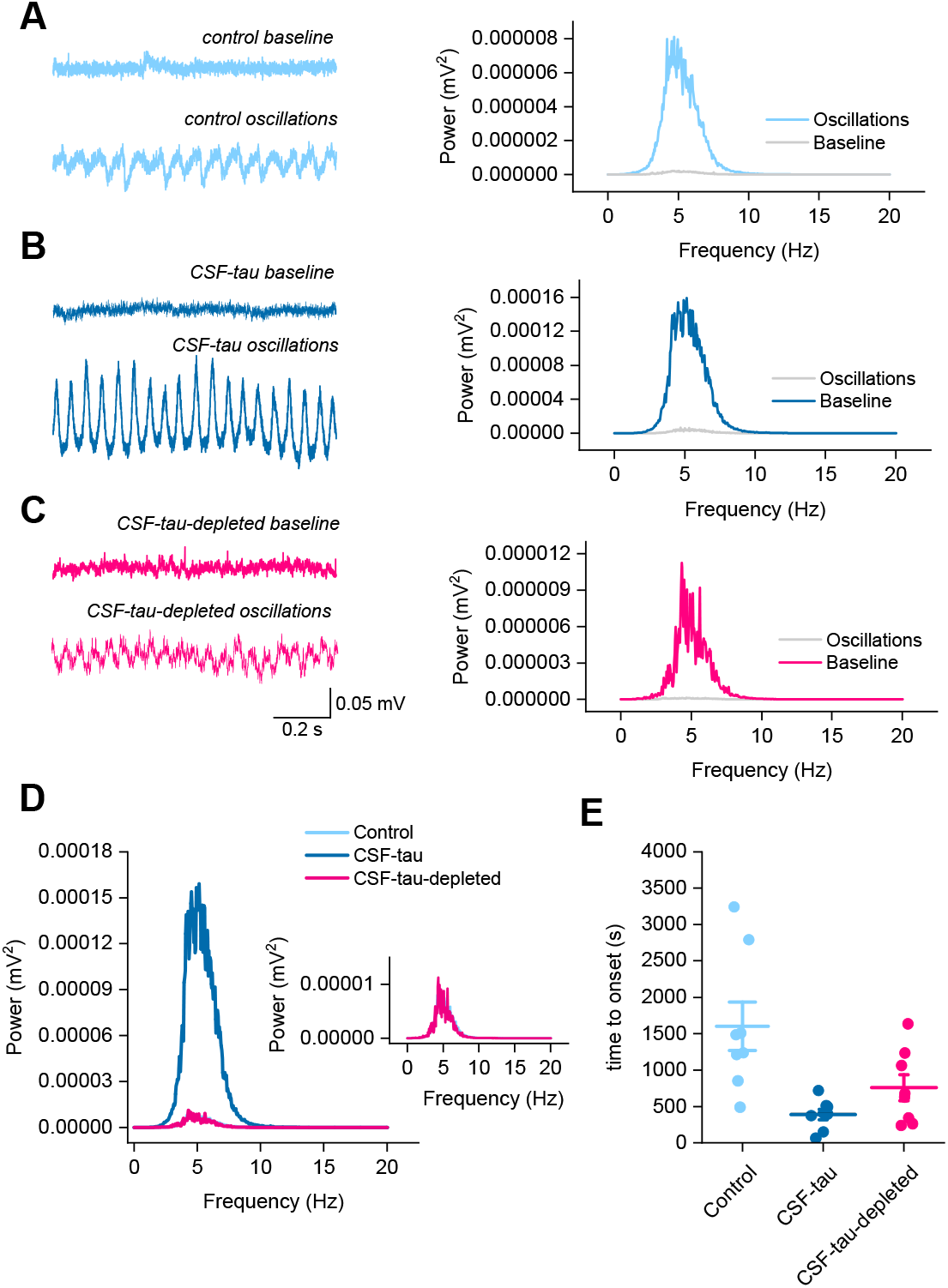
CSF-tau alters the generation and amplitude of theta oscillations. Theta oscillations in the CA3 region of the hippocampus were recorded in an interface chamber (see methods for details). Bath application of carbachol (50 μM) in this study induced robust, reliable theta oscillations across all three experimental conditions (**a, b, c**). **Left,** Representative examples of baseline and theta oscillations for each of the three conditions. **Right,** Power spectrum from the representative example confirming the oscillations to be in the theta range (4-7 Hz). **d,** mean power spectrums for each of the conditions. CSF-tau incubated slices had significantly stronger oscillatory power compared to control or CSF-tau-depleted incubated slices (P < 0.0001). Inset, higher magnification of control and CSF-tau-depleted traces. **e,** CSF-tau incubated slices also showed a significantly quicker onset of oscillatory activity relative to control (p = 0.005). Data is presented as Mean ± SEM.

We also found that the amplitude of mEPSCs was significantly enhanced in CSF-tau incubated slices *(Figure 4d).* Using a Kolmogorov-Smirnov test to compare the cumulative frequency of the pooled mEPSCs for each of the conditions, we found that distribution of mEPSC amplitudes in slices exposed to CSF-tau (n=800 pooled mEPSCs from 8 slices) was significantly different to that of control slices (n=600 mEPSCs from 6 slices; D = 0.1533, p <0.0001). Then, by comparing the mean mEPSC amplitudes for each slice across the two conditions, we further validated that slices incubated in CSF-tau showed increased amplitudes compared to control slices (mean amplitude in CSF was 22.5 ± 0.39 pA vs mean amplitude in control was 18.9 ± 0.33 pA; Mann-Whitney test p = 0.0426, *Figure 4e*).

Finally, to establish whether there were any changes to the kinetics of the mEPSCs, we then measured the rise and decay time for the average waveform from each recording. The rise time was measured (10-90%) and the decay time was obtained by fitting with a single exponential. There was no change to rise or decay kinetics between the two conditions.

### 3.5 CSF-tau alters the generation and maintenance of theta oscillations

We have found that CSF-tau markedly changes neuronal properties, synaptic transmission, and synaptic plasticity. We have now investigated how these changes can lead to alterations in emergent network activity. Hippocampal hyperexcitability has been identified in AD patients prior to the onset of cognitive symptoms and is associated with increased oscillatory power in theta bands relative to control or MCI groups [45, 64, 89, 90, 98, 109, 111]. Thus, changes in theta power are of interest as a potential predictor for the development of AD [68].

Bath application of the acetylcholine receptor agonist carbachol (50 μM) induced robust, reliable theta oscillations, as demonstrated by Wilcoxon tests comparing peak oscillatory amplitude and baseline activity for all three treatment groups *(Figure 4a-c).* In control slices pre-incubated with aCSF (Figure 4a), there was a significant difference between baseline power and oscillatory power (W = −138601, z = −2.833e-008, p < 0.0001). In slices pre-incubated with CSF-tau (Figure 4b), the difference between baseline power and oscillatory power was also significant (W = −138601, z = −2.830e-007, p < 0.0001). Finally, in slices pre-incubated with CSF-tau-depleted (*Figure 4c*), there was a significant difference between baseline power and oscillatory power (W = −138601, z =-2.214e-008, p < 0.0001). A Kruskal-Wallis test showed that the difference in peak oscillatory power between the three treatment groups was also significant *(Figure 4d;* H(2) = 166.2, p < 0.0001), with Dunn’s tests revealing a significant increase in theta power following CSF-tau treatment relative to aCSF control slices (p < 0.0001) and treatment with CSF-tau-depleted (p < 0.0001), whilst there was no significant difference between control and CSF-tau-depleted groups. The significant enhancement in the amplitude of hippocampal theta oscillations following incubation with CSF-tau aligns with previous published studies demonstrating that increased theta power is one of the first pathological changes observed in AD patients [14, 32, 36, 69, 89, 91]. Finally, the time to onset of oscillatory activity was also significantly different between groups (*Figure 4e;* H (2) = 10.09, p = 0.0065), with CSF-tau slices showing a significantly quicker onset of oscillations relative to control (p = 0.005) following carbachol application.

## 4. Discussion

CSF biomarkers for tau and phosphorylated-tau have proved to be reliable predictors/indicators of AD [18, 72]. Tau-based biomarkers can differentiate AD from unaffected individuals and other tauopathies with strong predictive values [72]. Moreover, tau-based biomarker levels change in a stepwise manner across the AD spectrum, allowing for more accurate disease staging, clinical diagnosis, as well as clinical trial recruitment and monitoring [72]. Pathological tau species in AD brains are post-translationally modified, including being phosphorylated and aggregated both in soluble (e.g., oligomers, protomers) and insoluble (neurofibrillary tangles) forms. We currently have high-performing biomarkers that reflect both phosphorylation (e.g., p-tau181) [74] and aggregation [62, 113] of tau in CSF. However, little is known about the toxic effects on brain function of these soluble tau forms in CSF, and this could shed new light on the novel biomarkers developed and applied clinically.

The pathological effects of soluble tau aggregates have been studied extensively in animal models [19, 43, 79, 102, 105]. These changes are normally induced by full-length soluble aggregated forms of tau, mostly produced recombinantly [6, 33, 59, 85]. However, recent work has suggested that tau aggregates produced *in vitro* do not recapitulate the structural features of authentic AD brain tau [132]. We therefore sought to investigate if CSF-tau, which contains high levels of aggregated and non-aggregated tau, often in truncated forms [31], modulates neuronal function.

To do this, we incubated acute mouse brain slices in biomarker-quantified tau-positive CSF pooled from multiple patients. As the goal for future studies is to scale down the volume of sample from pooled samples (5 ml in this study) to individual patient samples (~ 300 μl), to enable correlation with clinical data, we first determined the minimum volume of CSF-tau (diluted in aCSF) for which we could detect a change to neuronal function. We then performed a suite of electrophysiological tests to evaluate the effect on neuronal and network function. We demonstrate that CSF-tau markedly increases neuronal excitability, alters synaptic transmission and plasticity, and modifies the generation and maintenance of hippocampal theta oscillations.

### 4.1 Immunodepletion of CSF pool to remove tau species

A wide range of tau fragments can be found in CSF. These are mostly N-terminal to mid-domain tau fragments and provide good biomarkers for disease prognosis, diagnosis, and staging [18, 108]. For this reason, in this study we have used a combination of three antibodies to provide a robust removal of N-term and Mid-terminal tau species: Tau12 N-terminal [epitope = amino acids 6-18], HT7 mid-region [epitope = amino acids 159-163], and TauAB C-terminal antibody [epitope = amino acids 425-441]. To verify the removal of tau after the immuno-depletion, we used different assays that cover various longer (Simoa t-tau assay from Quanterix and p-tau181 [73] and p-tau231 [8] in house assays) and shorter (Lumipulse p-tau181 assay [80]) tau fragments, respectively. Although less abundant, recent studies have shown evidences of other tau fragments in CSF, including microtubule-binding region (MTBR)-containing species [62, 63]. In this initial study, we did not include antibodies specifically targeting the MTBR of tau. Therefore, we cannot guarantee the extent of removal of this region of the protein. However, we demonstrate significant reduction of total and phospho-tau and significant rescue of the functional effects on neuronal function and oscillations when tau is removed. Thus, the observed changes can be attributed to CSF tau. Additional studies using different antibody panels are nevertheless planned in order to further confirm these results and expand them. While we cannot be certain that all of the observed effects are due solely to tau, as it may exist as a complex with other proteins, we can conclude that, as immuno-depleting for tau abolished the effects, tau must play a central role in mediating the changes.

### 4.1 CSF-tau led to an increase in excitability and decrease in whole-cell conductance

We used both standard IV (with step current injections) and the DIVs (with naturalistic current injection) to extract neuronal parameters. In slices incubated with CSF-tau, both approaches for parameterisation measured a marked decrease in whole-cell conductance (rise in neuronal input resistance) and a depolarisation of the resting membrane potential leading to an increase in firing rate (a correlate of neuronal excitability). Obtaining comparable results using two independent methods for the extraction of neuronal parameters strengthens the robustness of this observation.

Depolarisation of the resting membrane potential and neuronal hyperexcitability are consistent with previously reported data in numerous tauopathy models [59, 105, 118, 128], for example in the rTg4510 mouse model, which expresses human tau variant P301L, where the pyramidal cells were depolarised by ~8 mV compared to WT littermates [105]. The increase in input resistance and firing rate, coupled with the observed depolarisation could result from the block of a standing (or leakage) conductance for example a two-pore-domain potassium (K2P) channels that contribute to maintaining the resting potential [84].

A comparable increase in neuronal hyperexcitability has also been observed in AD patients in individuals performing memory-encoding tasks [27, 39, 54]. This phenotype is observed in the early stages of disease, before pronounced cell loss and hypoactivity [12, 46]. Thus, our *in vitro* findings compliment the published literature both for animal models and for early AD when measured in patients [12, 46].

### 4.2 CSF-tau enhanced synaptic transmission and plasticity

Soluble tau aggregates have been shown to alter synaptic plasticity both in *in vitro* and *in vivo* models of tauopathy [1, 43, 102, 104]. Therefore, we next investigated if CSF-tau altered synaptic plasticity by measuring LTP. In slices incubated with CSF-tau, we observed enhanced input-output responses. If this was solely due to an increase in release probability, one might expect that there would be less paired pulse facilitation, however, this is not what we observed. Slices incubated with CSF-tau showed significantly enhanced paired pulse facilitation, suggesting that while there may be a change in release probability, there may also be changes in post synaptic receptor numbers and/ or calcium processing.

We then recorded AMPA receptor mediated mEPSCs. In slices incubated with CSF-tau, we observed significant increases in both the amplitude and frequency of mEPSCs. An increase in mEPSC frequency would suggest an increase in release probability, consistent with the increased input-output responses, which could result from an increase in the number of active zones or the number of docked vesicles [9, 97, 131]. The increase in mEPSC amplitude could reflect an increase in the number of postsynaptic receptors, altered vesicle size or an increase in the number of synapses if the mEPSCs are multiquantal [66, 83, 131]. It is also possible that synapses that were previously silent (exhibiting NMDA-receptor mediated synaptic transmission but lacking AMPA receptors) might be unsilenced (via insertion of AMPARs) to give more functional synapses [122]. Silent synapses, which are normally found at the tips of dendritic protrusions called filopodia, allow neurons to maintain more potential for plasticity and have recently been demonstrated to be more abundant in adult mice than once thought [122]. In the rTg4510 tauopathy mouse model, a loss of spine density in the cortex is reported over time and this results from a reduction in mushroom spines. However, in contrast, filopodia are increased [35, 120], suggesting that tau pathology could alter the number of silent synapses and this could contribute to changes in plasticity. In future studies, it would be interesting to examine the ratios of GLUR1 / GLUR2 and also to look at AMPA phosphorylation as phosphorylation of GluA1 at serine 831 and 845 are reported to play key roles in synaptic plasticity [2].

### 4.3 Alteration to the generation and maintenance of theta oscillations

Functional theta oscillations are dependent upon synchronised activity within the hippocampus, which in turn is mediated by pyramidal cells and interneurons [50]. Parvalbumin (PV)-expressing interneurons are suggested to be particularly critical in regulating pyramidal cell activity [5, 99, 123].

In this study, we have demonstrated that incubation of brain slices with CSF-tau markedly enhanced theta oscillations and increased excitatory pyramidal neuron excitability, and that both of these effects were abolished upon immunodepleting for tau. It is currently unclear whether the increase in excitability of pyramidal cells and changes in glutamatergic transmission are sufficient to produce the observed increases in theta power or whether changes in GABAergic inhibition also occurred. Our findings align with existing data in rodent models and AD patients; the deposition of tau in pyramidal cells and PV interneurons from 1 month of age in 3xTg mice was associated with elevated theta oscillations but no behavioural impairments [88]. The implication that network alterations occur early in the disease course is supported by the oscillatory slowing (an increased power in the theta band) observed in early AD patients accompanied by neuronal hypersynchronisation. Neuronal loss and a global discoordination of network activity then follows in later stages, alongside power alterations in other oscillatory bands such as alpha and delta [22, 41, 42, 52, 53, 76, 95, 96, 101, 106, 115, 116].

This leads to the question of what incubation with CSF-tau in this study might be doing to hippocampal neurons to mediate the elevation in the power of theta oscillations. A combination of experimental and simulated data has indicated that the early-stage AD hyperactivity underpinning oscillatory slowing could be due to pyramidal neuron hyperexcitability [24, 26, 47, 94, 114, 134] and/or reduced excitability of GABAergic PV interneurons and thus pyramidal cell disinhibition [4, 30, 57, 86, 94, 100, 117]. Support for the latter hypothesis of disinhibition also comes from the loss of inhibitory synapses in AD along with the pronounced GABAergic dysfunction observed in AD mouse models, including the model of APOE4, the principal genetic disease risk factor [23, 94]. Such findings provide a potential mechanism for the oscillatory disruption induced by CSF-tau incubation in this study and imply such network alterations to represent an element of the core, initial neuropathology of AD. Indeed, recent studies are now evaluating the therapeutic implementation of theta entrainment for AD patients [38, 81, 129, 130].

It has also been demonstrated that changes to theta oscillatory power – similar to what we observed herein – can be used to distinguish between prodromal AD and non-AD cases with cognitive decline [94]. The fact that our effects on oscillations agree with clinical human electrophysiological studies gives confidence that this is a method that could, in the future, be clinically useful as in compliment to EEG testing.

### 4.5 Conclusion

Incubation of acutely-isolated wild-type mouse hippocampal brain slices with small volumes of diluted human CSF-tau allowed us to evaluate effects on neuronal function from single cells through to network level effects. Comparison of the toxicity profiles of the same CSF samples, with and without immuno-depletion for tau, enabled a pioneering demonstration that CSF-tau potently modulates neuronal function. We demonstrate that CSF-tau mediates an increase in neuronal excitability in single cells. We observed, at the network level, increased input-output responses and enhanced paired-pulse facilitation as well as an increase in long-term potentiation. Finally, we show that CSF-tau modifies the generation and maintenance of hippocampal theta oscillations, which have important roles in learning and memory and are known to be altered in AD patients. Together, we describe a novel method for screening human CSF to understand functional effects on neuron and network activity, which could have far-reaching benefits in understanding pathological mechanisms of tauopathies, thus allowing the development of better targeted treatments.

## Declarations

### Ethics approval and consent to participate

CSF collection and processing were performed at the University of Gothenburg, Sweden, with local ethical approval, and the samples sent to the University of Warwick, UK, where all experiments were done as approved by the local Human Tissue Authority and Biomedical & Scientific Research Ethics Committees. All animal care and experimental procedures were reviewed and approved by the institutional animal welfare and ethical review body (AWERB) at the University of Warwick.

### Consent for publication

All authors have approved the manuscript for submission and that the contents have not been published or submitted elsewhere.

### Availability of data and material

The raw data generated and analysed for this manuscript are available from the corresponding author upon reasonable request.

### Authors’ contributions

Conceptualization: **EH, MJW, TKK, JB,** Methodology: **EH, MJW, TKK, JB,** Formal analysis, and Investigation: **EH, MJW, JB, EC, JLR, MO, AW, BM,** Writing - original draft preparation: **EH,** Writing - review and editing: **EH, MJW, TKK, JB, EC, JLR, MO, AW, BM, HZ, KB,** Resources: **EH, MJW, TKK, EC, JLR, MO, HZ, KB,** Supervision: **EH, MJW, TKK**

## Acknowledgements

We would like to thank Professor Bruno Frenguelli for providing an interface chamber for the recording of theta oscillations.

## Supplementary material

**Supplementary figure 1.**
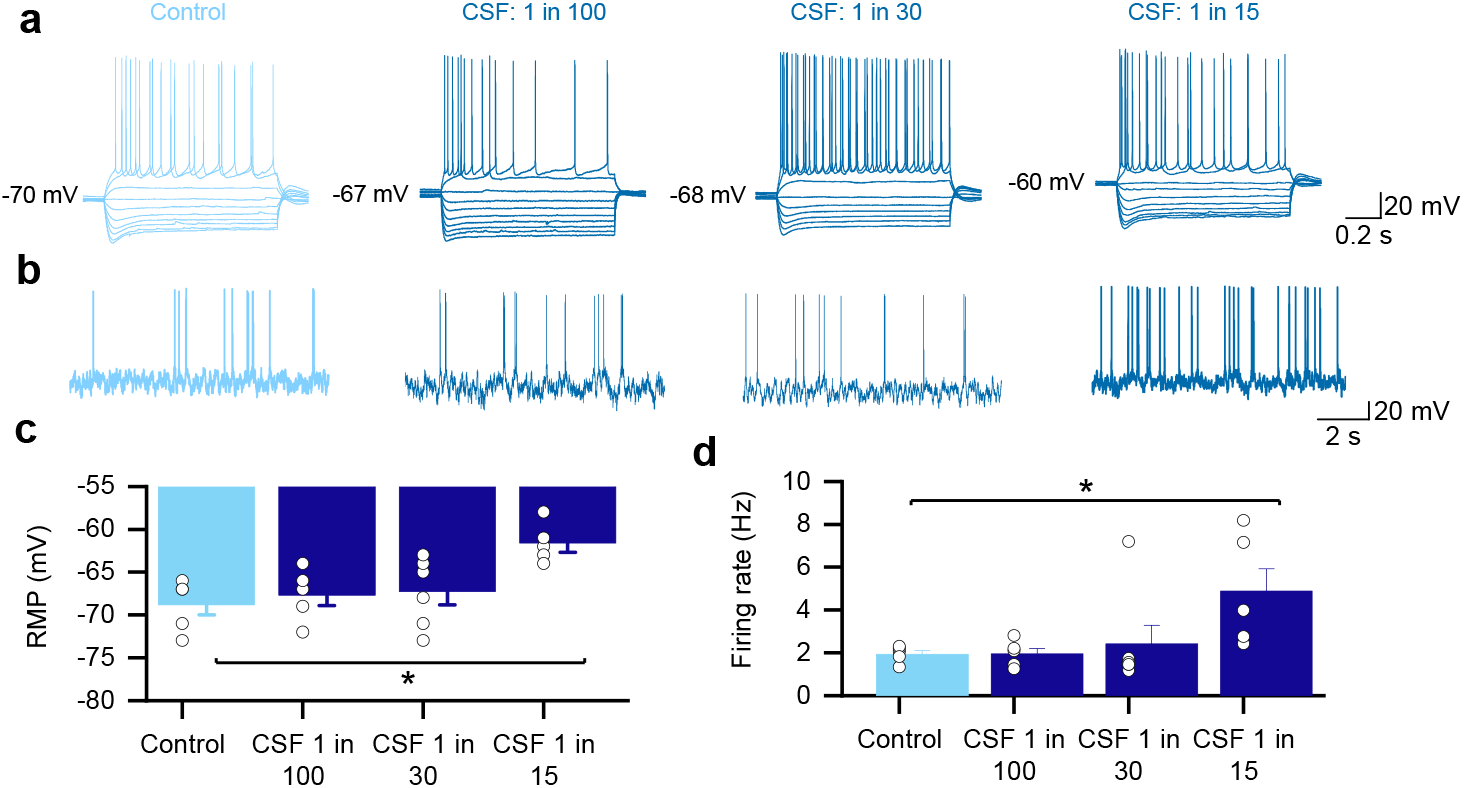
We diluted the CSF-tau sample with artificial CSF (aCSF) to reduce the volume of sample that was used for each experiment. As the end goal will be to use individual patient samples (much smaller volumes ~300 μl) to enable correlation with clinical data, we first sought to find the minimum volume (maximum dilution in aCSF) of CSF-tau, for which we could detect a change in neuronal function. Representative examples of standard current-voltage responses for slices that have been incubated in control aCSF vs different dilutions of CSF in aCSF (1:100, 1:30 and 1:15 dilutions). Using neuronal excitability, measured using whole-cell patch clamp recording from single CA1 pyramidal cells in the hippocampus as a readout. We found no significant changes to neuronal excitability with either 1:100 dilution or 1:30 dilution of CSF-tau, but a significant depolarisation (**a**; p = 0.0212) and increase in firing rate (**b**; p=0.0215) was observed with a 1:15 dilution (100 μl CSF in 1.5 ml aCSF) compared to control slices.

**Supplementary figure 2.**
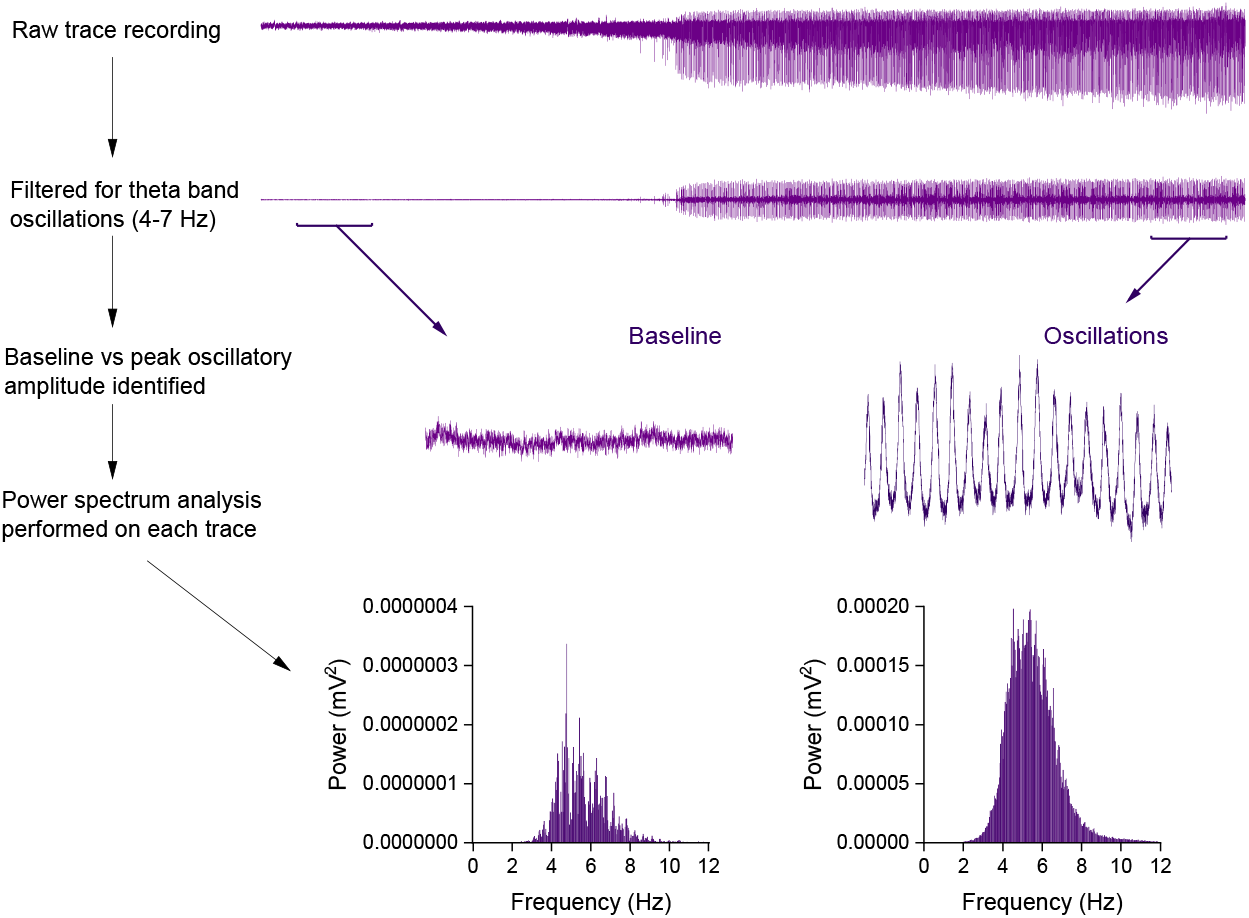
Protocol for analysis of hippocampal CA3 theta oscillations. Carbachol-elicited oscillations were characterised using power spectral density (PSD) analysis in Spike 2. PSD profiles of the field potential recordings filtered for the theta band (4 - 7 Hz) were generated by Fourier transform analysis (Hanning window, FFT size 2048, resolution 4.883 Hz) from each recording. The profiles were calculated from a 100 - 300 second section of the field potential trace displaying peak oscillatory activity, with the baseline power subtracted offline. Power spectrum analysis was performed to ascertain oscillatory power.

